# Annexin A4 senses membrane curvature in a density dependent manner

**DOI:** 10.1101/2021.11.02.466919

**Authors:** Christoffer D. Florentsen, Joshua A. Daniels, Guillermo Moreno-Pescador, Iliriana Qoqaj, Jesper Nylandsted, Poul Martin Bendix

## Abstract

Annexins (ANXs) are a family of peripheral membrane binding proteins which play a vital role in the maintenance and function of cellular membranes. These proteins are known to be part of the plasma membrane repair machinery where they are known to bind to negatively charged lipids in the cell membrane in a calcium dependent manner. The shape of the plasma membrane is known to be a regulator of protein density and thereby affects several biological functions of the cell such as exo-and endocytosis, cell motility, immune responses and also membrane repair. Membrane deformation and curvature sensing by proteins is a well described phenomenon which can assist in recruitment of specific proteins to certain regions in the cell and facilitate membrane bending. Following ruptures in the plasma membrane, calcium influx assist in the association of ANXs with the membrane around the ruptured area. Due to the expected increase in curvature at the damaged membrane site, it has been suggested that both membrane curvature and Ca^2+^ participate in the recruitment of ANXs. We have investigated the curvature sensing of ANXA4 in giant unilamellar vesicles (GUVs) by pulling high curvature membrane tethers from the vesicle surface using optical tweezers showing that ANXA4 recruitment increases with higher membrane curvature. We also describe an assay for determining protein density on the plasma membrane by utilizing ANXA4’s property as a calcium dependent membrane binding protein. This new assay allows us to investigate the effect of protein density on curvature sensing.

## Introduction

The ability of the cell to control the PM integrity is vital for cell survival and several proteins have been reported to be involved in the process of plasma membrane (PM) repair. In recent years, the protein family of ANXs has been suggested as being in the forefront of the PM’s defense system by upconcentrating in areas where the plasma membrane integrity has been compromised [1]. As several ANXs are known to bind to acidic phospholipids in a Ca^2+^ dependent manner, the influx of Ca^2+^ caused by PM rupture is expected to recruit ANXs which constitute an important part of the PM repair machinery [1].

The ANX protein family consist of 12 members, each with their own N-terminal sequence. The core consist of 4 highly conserved domains (8 for ANXA6) with Ca^2+^ binding sites, shaped into a disk-like structure with a slight bend[2]. Several members of the ANX family are known to interact with the PM when Ca^2+^ is present, thus reacting to influx of Ca^2+^ from the extracellular space.

We have previously investigated the recruitment of ANXA2, ANXA4 and ANXA5 to areas of high curvature in cell derived plasma membrane vesicles [3, 4] and the sorting of ANXs seems to be coupled to the ability of some ANXs to form trimers and counteracted by formation of extended immobile scaffolds on the membrane -a feature possessed by ANXA4 and ANXA5 [5, 6]. In this context, both curvature sensing and mobility of proteins may depend strongly on protein density and hence a method for determining the amount of protein recruited to the PM after damage which causes an influx of Ca^2+^, remains to be developed for these proteins. While the curvature sensing of ANXs in plasma membrane vesicles may provide insight into the behavior of the protein in a natural plasma membrane the observation may however include many different effects from interactions with other proteins and from the complex lipid composition. Moreover, the density of proteins on the membrane may play an important effect on both oligomerization and membrane curvature sensing as observed for other curvature sensing proteins, see Ref. [7]. Therefore, to elucidate the mechanism of membrane curvature sensing for specific annexin proteins an assay is needed which can be used to quantify the curvature mediated recruitment in a system where protein density, lipid composition and membrane curvature are known.

So far, no studies have evaluated the curvature sensing of ANXs in relation to its density at the PM, as the density of membrane bound ANX is not readily determined. To this end we have developed an assay which allows accurate quantification of the membrane bound protein density. The assay exploits the feature that annexin translocates from the vesicle lumen to the membrane upon addition of Ca^2+^. Since all ANX is recruited to the membrane, the density of membrane bound ANX can be determined by relating the lumen fluorescence intensity to vesicle geometry [7–9]. This allows us to quantify the bound density of ANX in each GUV and relate this to the membrane curvature sorting in membrane tubes extracted from the GUV using an optical trap [3, 4].

## Results and discussion

GUVs as a simplified model membrane system is a functional and well established assay used for investigation of protein organization on the PM and their interaction with membrane lipids. The simplicity of the system offers great control of the experimental factors as well as easy manufacturing and reproducibility [7, 10].

Here, we present a way of investigating the membrane curvature affinity of ANXA4 in a GUV system containing acidic phosphatidylserine (DOPS) and neutral phosphatidylcholine (DOPC) lipids. We use GUVs composed of of 79% DOPC, 20% DOPS and 1% biotin-X DHPE lipids and labeled using trace amounts of DiD (see Figure 1) to investigate the curvature affinity of encapsulated recombinant ANXA4, tagged with superfolder Green Fluorescent Protein (sfGFP). The formation of these vesicles is schematically depicted in Figure 1 together with representative confocal images of the GUVs.

**Figure 1:**
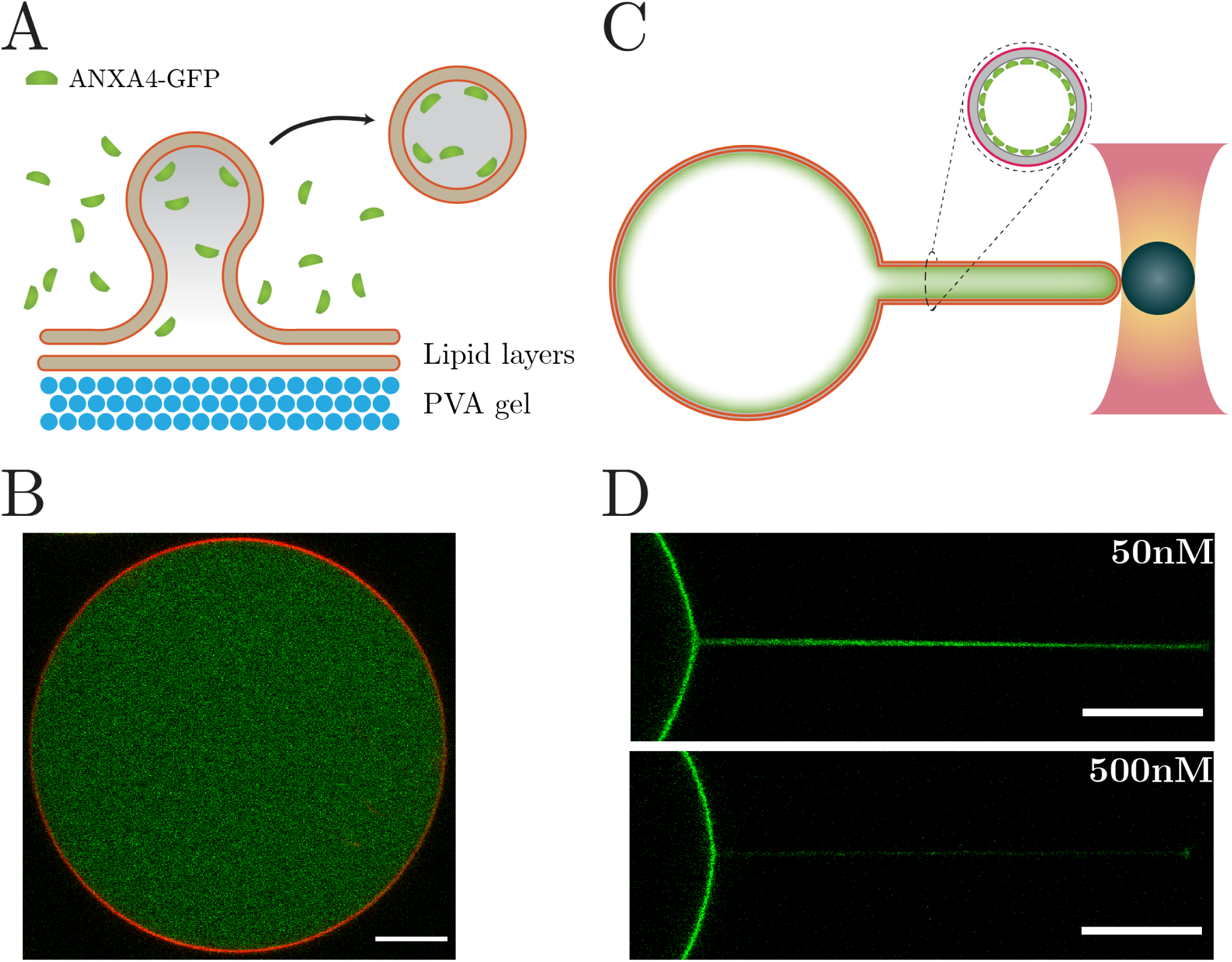
(A) Schematic illustration showing the production of GUVs using PVA coated glass cover slips. The membrane is labeled using vybrant DiD (red). (B) Confocal image (overlay) showing that the GUVs contain recombinant ANXA4-sfGFP encapsulated inside the GUV during hydration from the lipid film on the PVA gel. The exterior protein is washed away prior to usage of the vesicles. the membrane is shown in red and the proteins are shown in green. (C) Schematic showing the pulling of a tether from the GUV membrane using optical tweezers and a polystyrene bead. The up-concentration of ANXs is shown inside the resulting nanotube. The vesicles have been exposed to Ca^2+^ prior to pulling and hence the ANXs are depicted as membrane bound. (D) Confocal images showing nanotubes pulled from GUVs with membrane bound ANXA4-sfGFP (green). The vesicles produced with a 50nM concentration show a higher sorting than the vesicles produced using 500nM concentration of ANXA4-sfGFP. All images have been contrast enhanced with FIJI. All scalebars are 10 *µ*m.

### Encapsulation of proteins in GUVs

The GUVs were prepared using the protocol by Weinberger et al. [11], as described in the Experimental section. Briefly, a 2 mM lipid solution was mixed in chloroform stabilized with ethanol. After spreading a PVA solution on a coverslip and letting it dry, the lipid mixture was added and dried using nitrogen gas followed by drying in a vacuum chamber. ANXA4-sfGFP containing Growing Buffer (see Experimental Section) was added and the vesicles formed during 90 minutes. The vesicles were harvested by pipetting the solution into an Eppendorf tube followed by 10 minutes of centrifugation at 600 rcf and 13°C. The vesicle rich pellet was isolated and used for experiments. The GUVs were placed on a glass surface for observation in a confocal microscope. The use of C-terminal tags has previously been reported without any notions on the membrane binding properties, localization, folding or orientation for ANXA4, ANXA5 and ANXA6[3, 12]. Hence, we do not expect any influence of the sfGFP tag on our results.

For curvature sensing experiments, nanotubes were pulled from the GUVs by using an optical trap. This allows for measurements of the mobility and partitioning of the fluorescently labeled proteins in a controlled manner inside the nanotube which has significantly higher curvature than the vesicle as depicted in Figure 1C.

### Protein concentrations are high inside the nanotubes

To investigate how the sensing abilities of ANXA4 is affected by the concentration (and membrane coverage) of proteins, we present a simple method for calculating the sfGFP density on a vesicle membrane. Note that for this procedure any sfGFP tagged ANX can be used which can translocate all the encapsulated sfGFP-tagged protein from the lumen to the membrane upon addition of Ca^2+^. The method comprises four steps:

i. A calibration curve is measured in solution which relates ANXA-sfGFP fluorescence intensity to the concentration
ii. ANXA-sfGFP is encapsulated in an un-bound state within the GUV lumen in absence of Ca^2+^
iii. The concentration of ANXA-sfGFP within the GUV is found from the sfGFP intensity by using the calibration curve relating intensity and concentration from (i)
iv. By addition of Ca^2+^ to the exterior medium residual leakage of Ca^2+^ occurs into the GUV and consequently all the ANXA-sfGFP translocates from the lumen to the interior membrane of the GUV. From the geometry of the GUV and the internal concentration (determined in (iii)), the protein density on the membrane is quantified (see Materials and Methods).

For the quantification of the concentration dependent fluorescence intensities of sfGFP, an observation buffer containing glucose, Tris and NaCl was prepared and recombinant sfGFP-ANXA4 was added (see Experimental section). The solution was added to a cover slide for observation in a confocal microscope. By measuring the average pixel intensity at different ANXA4-sfGFP concentrations in solution, an intensity calibration curve was obtained as shown in Supplementary Figure S1A. As expected there is a clear linear relation between the protein concentration and the average pixel intensity expressed by Eq. (1). As the background signal has been removed from the intensity values, the *k* constant equals 0.

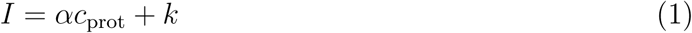

The values of *α* are shown in Supplementary Table 1. It is evident that similar linear relations are found for all laser powers (see Supplementary Figure 1), which enables extraction of *α* values for all laser powers in our current setup.

To determine the internal protein concentration in GUVs it is sufficient to known the protein concentration of the swelling buffer for GUV preparation and the encapsulation efficiency during formation of the GUVs. Efficient encapsulation of ANXA4-sfGFP during growth is essential. By swelling the GUVs in a medium containing recombinant ANXA4-sfGFP and the membrane dye Vybrant DiD, incorporation of protein into the lumen was achieved, see Figure 1 and Figure 2.

**Figure 2:**
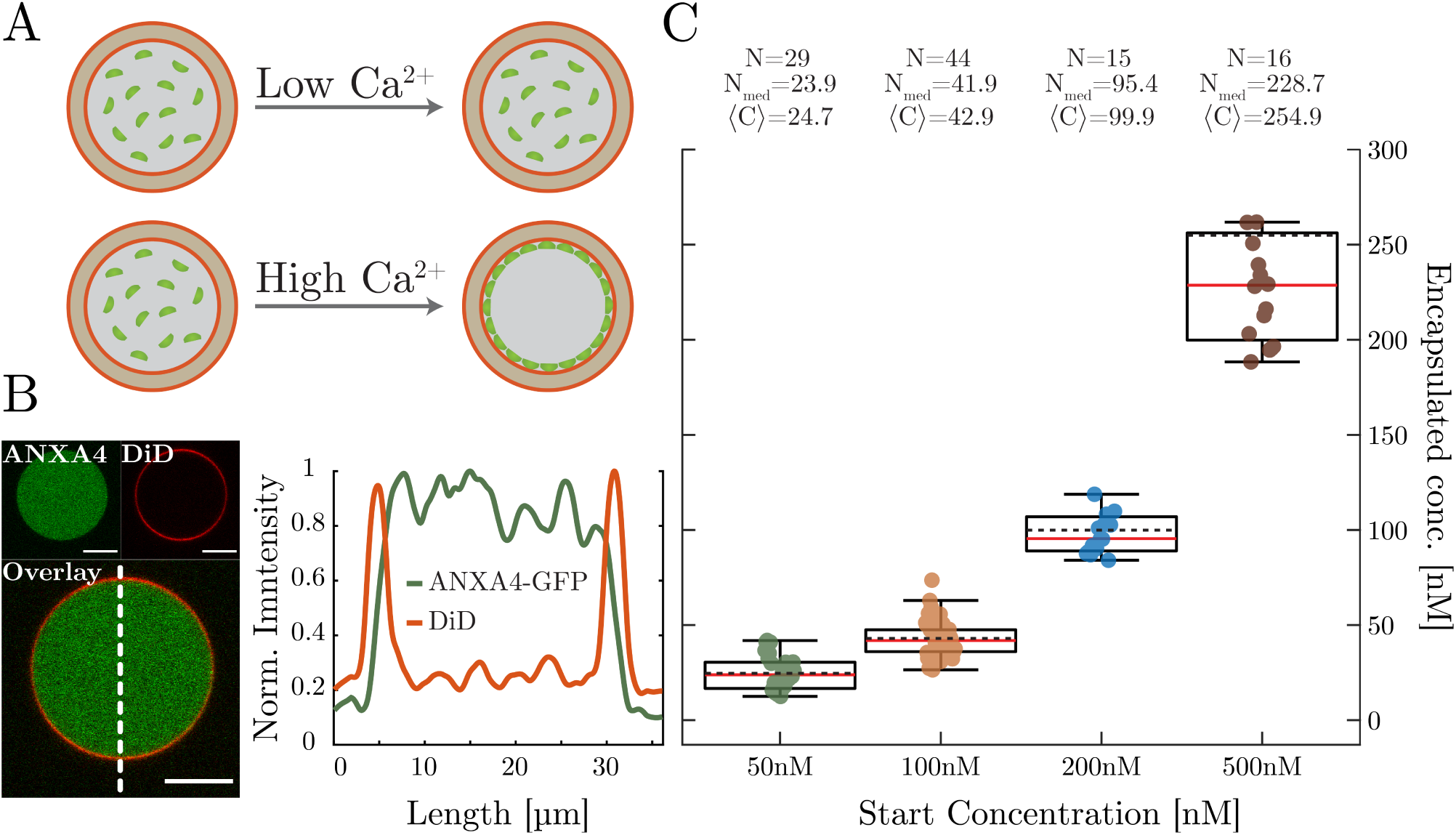
(A) Schematic showing the localization of ANXA4-sfGFP before and after addition of Ca^2+^. At low Ca^2+^ concentration, the ANXA4-sfGFP remains unbound and in solution. As the concentration of Ca^2+^ increases, the proteins start to bind to the membrane. (B) Left: Confocal images showing ANXA4-sfGFP and membrane label DiD as well as an overlay with the location of the lineprofile indicated by the dotted line. Right: Normalized pixel intensity of encapsulated ANXA4-sfGFP after addition of Ca^2+^. Red: membrane DiD, green: ANXA4-sfGFP. Scale bars are 10*µ*m and the images have been contrast enhanced in Fiji[13]. (C) Encapsulated concentrations for four different starting concentrations of ANXA4-sfGFP. The red lines denote the median and the black dotted lines denote the mean. It is clear that the encapsulation efficiency is found to be ∼50% of the starting concentration. Please note that outliers have been removed for simplicity. The plot containing the outliers is found in Supplementary Figure S2.

ANXA4 is known to bind to negatively charged membranes, but only in the presence of Ca^2+^. The GUV preparation buffer (i.e growing buffer) did not contain any Ca^2+^, and the ANXA4-sfGFP is expected to stay inside the lumen. Excess non-encapsulated ANXA4-sfGFP is removed by centrifugation.The unbound, internalized, ANXA4-sfGFP was imaged in a confocal microscope and compared to the intensities of a reference solution containing a known concentration of recombinant ANXA4-sfGFP and the intensity ratio was used for determining the encapsulation efficiency (EE).

The average encapsulated concentration, C, for four starting concentrations of recombinant ANXA4-sfGFP is shown in Figure 2C. From the data in Figure 2C the average encapsulation efficiency (EE) can be found.

From the data in Figure 2C it is evident that the average concentration found inside the vesicles is approximately half the concentration used in the growth medium. Similar EE values were also obtained by Weinberger et al. [11]. Following determination of the EE, GUVs are introduced to a buffer containing Ca^2+^. The influx of Ca^2+^ facilitates binding of all ANXA4-sfGFP to the inner leaflet of the GUV membrane (an example is shown in Figure 3). With the knowledge of ⟨ C ⟩, the surface density of proteins in any give vesicle can be determined if all optical settings are kept constant. Hence, the surface density inside a nanotube can be compared to the surface density on the vesicle membrane. This allows quantification of the effect of protein density on membrane curvature sensing.

**Figure 3:**
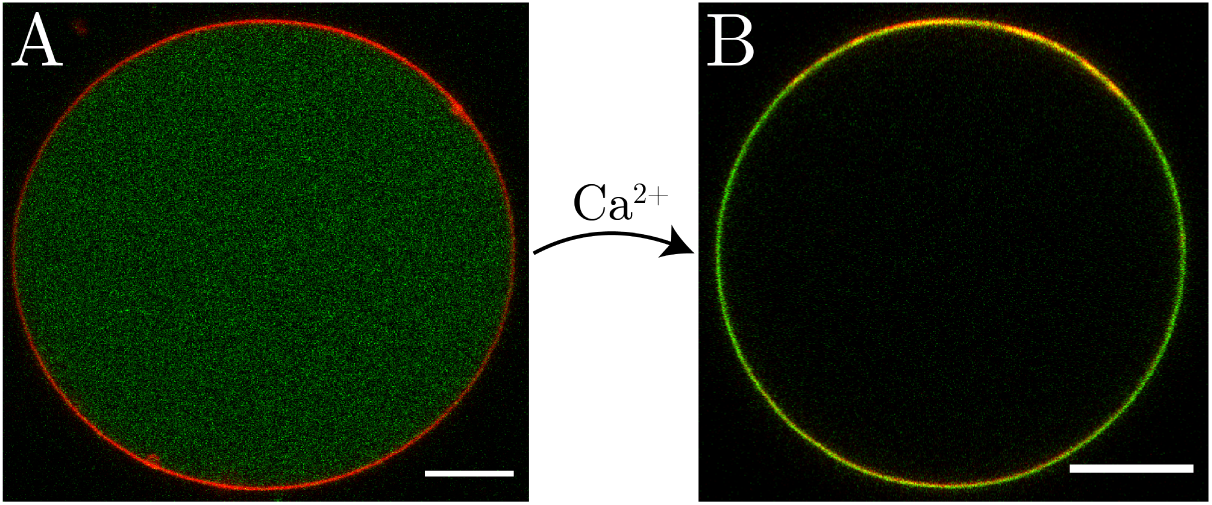
Surface density of proteins can be calculated knowing the size of the vesicle and the intensity of the proteins on the membrane at specific microscope settings. By addition of Ca^2+^, all the ANXA4-sfGFP becomes membrane bound and the surface density is readily determined from the vesicle size and the encapsulation efficiency. Scale bars are 10 *µ*m. The images have been contrast enhanced.

### ANXA4 senses negative membrane curvature

We have previously shown that ANXA4 and ANXA5 senses negative curvature in the presence of Ca^2+^ in giant PM vesicles (GPMVs)[3, 4]. Here, we investigate the membrane preference of ANXA4 in a simple lipid environment existing of two lipid species, DOPC and DOPS, and a trace amount of lipid dye Vybrant DiD as a reference marker for the membrane. We take advantage of the fact that the fluorescent dye does not have a curvature preference[14], thus being a reliable reference for the membrane area and hence its intensity scales with the radius of the tube [15]. Recruitment of ANXA4-sfGFP to the membrane was detected with confocal imaging. The density of proteins inside the nanotube can be determined relative to that on the vesicle membrane according to Eq. (2). Here, the intensity of the two fluorophores on the vesicle membrane (*I*_mem_) and the intensity measured inside the nanotube (*I*_vesicle_) are related and the sorting *S* is determined as:

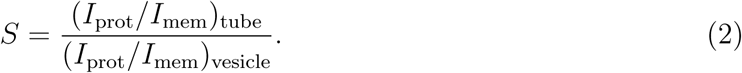

Following GUV formation and ANXA4-sfGFP encapsulation, membrane tethers were pulled from the vesicles and the sorting was quantified. The sorting values obtained from GUVs with ANXA4-sfGFP, are shown in Figure 4B. ANXA4 binds preferentially to the high curvature areas inside the nanotubes. Nearly all measured sorting values are larger than *S* = 1 with some showing a 6-fold increase in nanotube preference. All the nanotubes pulled from the GUVs are below the optical resolution of the confocal microscope (∼ 200 nm). It has been demonstrated that the common range of radii for nanotubes extracted from cells and vesicles lie in the window between 20-240 nm[7, 14, 16, 17]. As a micropipette aspiration assay has not been used in the current study, the specific absolute radial dependency of the sorting is not quantified. However, since the diameter of the tube scales with the intensity of the membrane dye we can still plot the sorting data versus a relative diameter which scales linearly with the physical tube diameter, see inset of Figure 4B.

**Figure 4:**
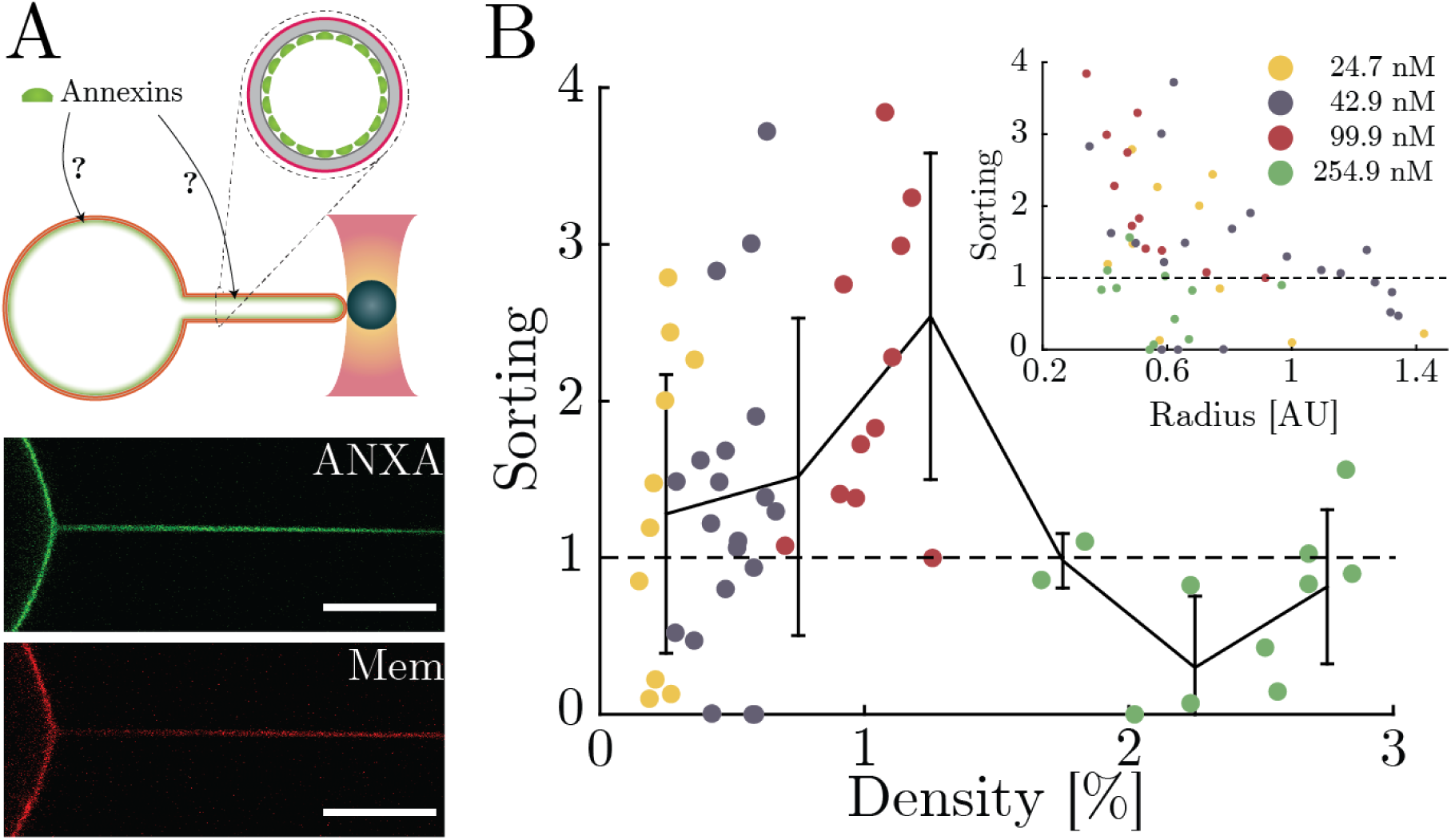
Membrane curvature sensing by ANXA4-sfGFP in GUVs. The Sorting value indicates the density of protein in the highly curved nanotube relative to the density on the nearly flat vesicle membrane calculated from Eq. (2). (A) Schematic showing a pulled tube with the two fluorescence channels shown below (green: ANXA4-sfGFP, red: DiD). The images have been contrast enhanced. The scale bars are 10*µ*m (B) Sorting data from four different starting concentrations of ANXA-sfGFP. The sorting is plotted as a function of protein density on the membrane. The solid line corresponds to the binned data median together with the interquartile range. The inset shows how sorting decays with increasing tube thickness. The x-axis scales linearly with tube radius and is given in arbitrary units.

### Sorting versus density

As the surface density is dependent on the radius of the spherical vesicle, the actual surface density may vary significantly depending on individual vesicle size. By performing single vesicle analysis using the vesicle radius, the surface density can be described in terms of the percentage of surface covered in each vesicle, see Materials and Methods. In the inset in Figure 4B, sorting is plotted both as a function of surface density and relative tube thickness. Surface densities of 0.25-3.5% are obtained for vesicles formed with ANXA4-sfGFP at starting concentrations of 50, 100, 200 or 500nM. A sorting maximum can be observed at densities of 1%.

Whereas previous sorting studies of I-BAR domain proteins have shown an inverse relation between surface density and sorting [7], our results indicate that curvature sorting is highest at intermediate densities of ANXA4 centering around 1%. This observation could be related to the ANXA4 trimer-formation, as ANXA4 sorting has been reported to be impaired for trimer-deficient ANXA4 mutants[3, 4]. As the surface density decreases, the ANXA4 proteins could have a lower trimer association rate, thus explaining the decrease in sorting for the lowest densities. At higher densities ANXA4 is known to form connected protein clusters which naturally will reduce the protein sorting.

I-BARs were additionally found to exhibit a maximum in sorting at a certain membrane curvature [7]. We were not able to detect such an optimal radius for sorting since it would require aspiration of the GUVs to obtain smaller tube radii. Notably, it was shown theoretically that a ANXA4 does optimally sense membrane curvatures corresponding to tube diameter of ∼40nm [4].

The results in Figure 4 show that ANXA4 senses membrane curvature with a preference for highly curved regions, as the tendency of measuring higher values of sorting increases for decreasing radii. The scattering in the data suggests that an underlying mechanism of ANXA4 membrane curvature sensing needs to be met before sorting happens. Since sorting of ANXA4 seems to be dependent on both tube radius and density further experiments are needed where tube radius is controlled by aspiration using a micropipette and protein density is quantified for each vesicle. Both density and tube radius could well affect the ability of the protein to form trimers and larger protein assemblies which can explain the scattering observed in our data [4].

### Positive Membrane Curvature

As the membrane curvature inside a nanotube is negative, we made control experiments determining if the ANXA4 would disfavour positive membrane curvature. These were carried out by extracting nanotubes from GUVs and by adding recombinant ANXA4-sfGFP to the external medium containing calcium and the DiD stained GUVs. Hence, the protein is only able to bind to the outer positively curved surface on the nanotube. Interestingly, we found that ANXA4-sfGFP is only a sensor for negative membrane curvature which was evident from the undetectable protein signal from the nanotube when ANXA4-sfGFP is added to the external medium (see Supplementary Figure 3). This suggests a lower preference of ANXA4 for positively curved membrane areas than for the “flat” membrane on the vesicle.

## Discussion

The membrane curvature sensing probed from the interior surface inside membrane nanotubes, extracted from GUVs, offers a great system to explore the dynamics of ANXA4 and measure curvature sorting in presence of a known density of proteins without any interfering effects from other proteins (Figure 4B, insert). The curvature preference is dependent on the density of proteins on the vesicle membrane, which is in good agreement with previously reported findings [7, 18]. The mechanism behind membrane curvature sensing by ANXA4 could be linked to the convex shaped ANXA4 core domain which binds to the membrane as well as the trimerization of the protein [4, 19]. Molecular curvatures are well known to facilitate curvature sensing and curvature generation[20]. ANXA4 has been shown to generate membrane curvatures in supported lipid bilayers [12] and ANXA4 was found to sense negative membrane curvatures in nanotubes extracted from giant plasma membrane vesicles (GPMVs) [4]. The use of GPMVs in Ref. [4] comprises a complex system with a crowded environment consisting of native plasma membrane proteins which could be competing for the same membrane binding sites as ANXA4. Hence, together with the results here, carried out with model membranes, ANXA4 can be concluded to both senses negative membrane curvatures in absence and presence of other proteins. The dependence of the sorting on the density could be linked to the ability of the protein to form trimers. The curvature sensing ability was found to depend on the trimerization of ANXA4 in Ref. [4], but at high densities we would expect a decrease in the sensing due to large scale oligomerization often observed for ANXA4 and hence decreased mobility.

As mentioned the trimerization ability of ANXs have been shown to be as a key factor during membrane repair [12, 21] and has been shown to be important for membrane curvature sensing of ANXA4 and ANXA5[3, 4]. In Ref. [3] it was observed that ANXA5 (as opposed to ANXA2 which does not form trimers) senses negative membrane curvature in GPMVs. We expect ANXA4 to form trimers in our experiments, however, the curvature sensing ability of the trimer versus monomer needs to be tested with a recombinant mutant of ANXA4 which is not able to form monomers.

It has been shown that curvature sensing proteins have a tendency to sense curvature of specific radii [7]. This phenomenon is also observed for ANXA4. Simulations have suggested that the trimeric form of ANXA4 has an optimal curvature range in which sensing is favorable [4]. To assess the full spectrum of curvature affinity of ANXAs aspiration of the GUVs has to be carried out in future studies.

## Conclusion

We have here quantified the curvature sorting of ANXA4 in lipid nanotubes and correlated the recruitment to density using a novel assay for measuring protein density. The assay for measuring protein density is based on GUVs encapsulating proteins which bind to membranes in presence of calcium. This method can easily be extended to any fluorophore-conjugated proteins.

Finally, we have shown that ANXA4 proteins exhibit a higher preference for highly curved membranes and exhibit stronger curvature sensing for lower protein densities.

## Materials and Methods

### Reagents

1,2-dioleoyl-sn-glycero-3-phosphocholine 18:1 (*δ*9-Cis)PC (DOPC) and 1,2-dioleoyl-sn-glycero-3-phospho-L-serine (sodium salt) 18:1 PS (DOPS) was purchased from Sigma-Aldrich. N-((6-biotinoyl)amino)hexanoyl)-1,2-dihexadecanoyl-sn-glycero-3-phosphoethanoamine, triethylammonium salt (biotin-x-DHPE) was purchased from Thermo Fisher Scientific. Polyvinyl alcohol (PVA) and *β*-casein from bovine milk was purchased from Sigma-Aldrich. Vybrant DiD cell-labelling solution (#V22887) was obtained from Thermo Fisher Scientific. Streptavidin coated microbeads (#CP01006) with a mean diameter of 4.95 *µ*m were purchased from Bangs Laboratories, Inc. Recombinant ANXA4-sfGFP was provided by Jesper Nylandsted and Theresa Louise Boye from The Danish Cancer Society.

### GUV Formation

The composition of the GUV were as follows: 79.2% DOPC, 20% DOPS, 0.6% DHPE-x-Biotin and 0.2% DiD. The lipids and the dye was mixed at a concentration of 2 mM in chloroform. For the growing buffer, we used 70 mM NaCl (sterile filtered through a 0.2 *µ*m filter), 25 mM Tris (pH 7.4), and 80 mM Sucrose. For the observation buffer, we used 70 mM NaCl (sterile filtered through a 0.2 *µ*m filter), 50 mM Tris (pH 7.4), and 55 mM Glucose. An observation buffer with 3mM CaCl_2_ was used for sample analysis of membrane bound ANXA4.

Polyvinyl alcohol (PVA) gel 5% (w/w) was placed in a 60^*°*^C laboratory heating oven for 30 minutes. 90 *µ*L PVA gel was added to a plasma cleaned cover glass and dried at 50^*°*^C for 50 minutes. 30 *µ*L of lipid solution was added to the PVA coated cover glass and dried with N_2_ gas followed by drying it in vacuum for 2 hours.

The cover glass was placed in a teflon chamber. The protein solution was diluted in growing buffer to obtain the desired concentrations. The buffer protein solution was added to the cover glass chamber (V_total_ = 300*µ*L). The chamber was covered to prevent light and air exposure and set to form GUVs for 1 hour. The vesicle containing solution was harvested and placed in the observation buffer (1 mL). The solution was centrifuged (600rcf for 10 minutes at 13^*°*^C) and 1mL of the supernatant was removed.

### Sample preparation

Vacuum grease was applied onto a cover glass #1.5 (24×40mm cleaned in 70% ethanol) to form rectangular 1.5×1.5 cm^2^ grease chambers. 50mg/L *β*-casein was added to the chamber to passivate the cover glass surface for 10 seconds and the chamber was subsequently washed 3 times with observation buffer. For samples without CaCl_2_ 40-100*µ*L of the GUV solution was added to the grease chamber. For samples with CaCl_2_ 16*µ*L GUV solution was mixed with 4*µ*L of microbeads and 180*µ*L observation buffer with 3mM CaCl_2_. 40-100*µ*L of the mixture was added to the grease chamber.

### Calibration curve measurement

A cover glass #1 (25mm ø) in a teflon chamber was passivated with 5g/L *β*-casein for 2 hours and washed 3 times with observation buffer. protein solution at the desired concentration was added to the chamber.

### Data analysis and density calculations

All data analysis was performed using Matlab (The Mathworks, Inc., Natick, MA). Custom-made scripts was used for analysis of the sorting ratios and the encapsulation efficiency. Briefly, the vesicle to nanotube protein ratios were quantified and a membrane dye signal (DiD) was used as reference. Bleaching was accounted for by normalizing to the signal from the vesicle membrane. All Matlab scripts are available upon request.

The protein density *σ* was calculated from the known area (*A*_*GUV*_) and volume (*V*_*GUV*_) of the GUV and from the protein concentration inside the vesicle, *c*_*ANXA*_, using the following equation:

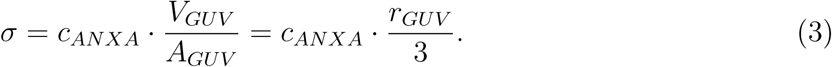

Subsequently, we calculate the area fraction, *ϕ* occupied by the protein as *ϕ* = *σ A*_*ANXA*4_. The area occupied by one ANXA4 protein was calculated in Ref.[4] to be *A*_*ANXA*4_ = 36nm^2^.

## Acknowledgments

This work is financially supported by Danish Council for Independent Research, Natural Sciences (DFF-4181-00196), by a Novo Nordisk Foundation Interdisciplinary Synergy Program 2018 (NNF18OC0034936), by the Lundbeck Foundation (R218-2016-534) and by the Danish National Research Foundation (DNRF116). The authors thank Anne Sofie Busk Heitmann (Danish Cancer Society Research Center) for recombinant protein production.

## References

1. Boye, T. L. & Nylandsted, J. Annexins in plasma membrane repair. Biological Chemistry 397, 961–969 (Oct. 2016).

2. Lizarbe, M. A., Barrasa, J. I., Olmo, N., Gavilanes, F. & Turnay, J. Annexin-phospholipid interactions. Functional implications. International Journal of Molecular Sciences 14, 2652–2683 (Jan. 2013).

3. Moreno-Pescador, G. et al. Curvature- and Phase-Induced Protein Sorting Quantified in Transfected Cell-Derived Giant Vesicles. ACS Nano 13, 6689–6701 (June 2019).

4. Florentsen, C. D. et al. Annexin A4 trimers are recruited by high membrane curvatures in giant plasma membrane vesicles. Soft matter 14, 2652 (Aug. 2020).

5. Newman, R. H., Leonard, K. & Crumpton, M. J. 2D crystal forms of annexin IV on lipid monolayers. Febs Letters 279, 21–24 (Feb. 1991).

6. Bouter, A. et al. Annexin-A5 assembled into two-dimensional arrays promotes cell membrane repair. Nature Communications 2, 274–9 (Apr. 2011).

7. Prévost, C. et al. IRSp53 senses negative membrane curvature and phase separates along membrane tubules. Nature Communications 6, 8529 (Oct. 2015).

8. Aimon, S. et al. Functional Reconstitution of a Voltage-Gated Potassium Channel in Giant Unilamellar Vesicles. PLoS ONE 6, e25529 (2011).

9. De Franceschi, N. et al. The ESCRT protein CHMP2B acts as a diffusion barrier on reconstituted membrane necks. Journal of Cell Science 132, jcs223644–23 (Aug. 2018).

10. Walde, P., Cosentino, K., Engel, H. & Stano, P. Giant Vesicles: Preparations and Applications. ChemBioChem 11, 848–865 (Mar. 2010).

11. Weinberger, A. et al. Gel-assisted formation of giant unilamellar vesicles. Biophysical Journal 105, 154–164 (July 2013).

12. Boye, T. L. et al. Annexin A4 and A6 induce membrane curvature and constriction during cell membrane repair. Nature Communications 8, 1623 (Nov. 2017).

13. Schindelin, J. et al. Fiji: an open-source platform for biological-image analysis. Nature Publishing Group 9, 676–682 (July 2012).

14. Rosholm, K. R. et al. Membrane curvature regulates ligand-specific membrane sorting of GPCRs in living cells. Nature Chemical Biology 13, 724–729 (2017).

15. Prévost, C., Tsai, F.-C., Bassereau, P. & Simunovic, M. Pulling Membrane Nanotubes from Giant Unilamellar Vesicles. Journal of Visualized Experiments, 1–9 (2017).

16. Ramesh, P. et al. FBAR Syndapin 1 recognizes and stabilizes highly curved tubular membranes in a concentration dependent manner. Scientific Reports 3, 1565 (Dec. 2013).

17. Tian, A. & Baumgart, T. Sorting of Lipids and Proteins in Membrane Curvature Gradients. Biophysj 96, 2676–2688 (Apr. 2009).

18. Sorre, B. et al. Nature of curvature coupling of amphiphysin with membranes depends on its bound density. Proceedings of the National Academy of Sciences 109, 173–178 (Jan. 2012).

19. Moss, S. E. & Morgan, R. O. The annexins. Genome Biology 5, 219–8 (2004).

20. McMahon, H. T. & Boucrot, E. Membrane curvature at a glance. Journal of Cell Science 128, 1065–1070 (Mar. 2015).

21. Lin, Y.-C., Chipot, C. & Scheuring, S. Annexin-V stabilizes membrane defects by inducing lipid phase transition. Nature Communications, 1–13 (Jan. 2020).

