## Supplementary Information for "Annexin A4 senses membrane curvature in a density dependent manner"

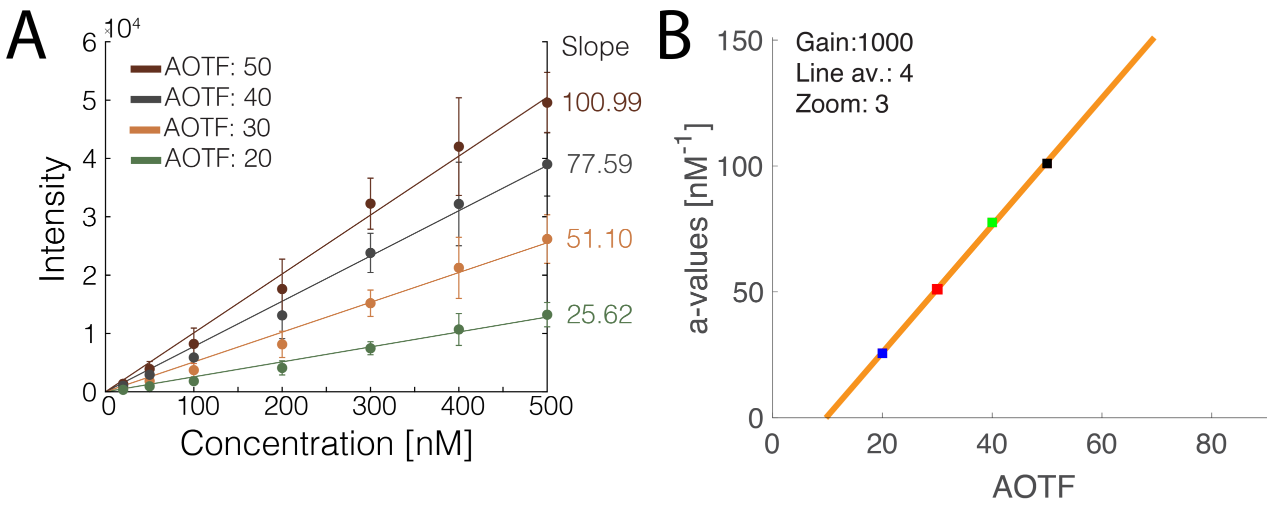


**Fig. S1.** (A) Calibration curves for ANXA5-sfGFP using zoom 3, line average 4 and gain 1000. The plotted intensities are compiled from three experiments. The background has been removed by imaging without ANXA5-sfGFP in the sample. Hence, the fit is set to cross through the point (0,0). (B) Plot of the slopes from the fitting in (A). The fit is linear allowing a determination of the slope at any AOTF, given the settings are kept constant. The resulting fit is given by the equation $a_{\text{value}} = 2.5259 P_{\text{AOTF}} - 24.5769$

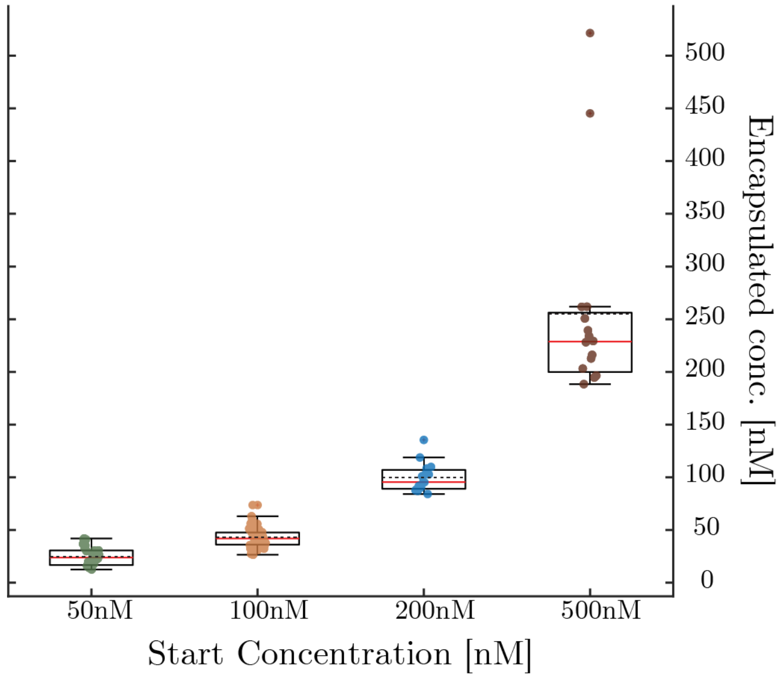


**Fig. S2.** Plot of encapsulation concentrations containing the outliers that have been removed from the plot in Figure 2

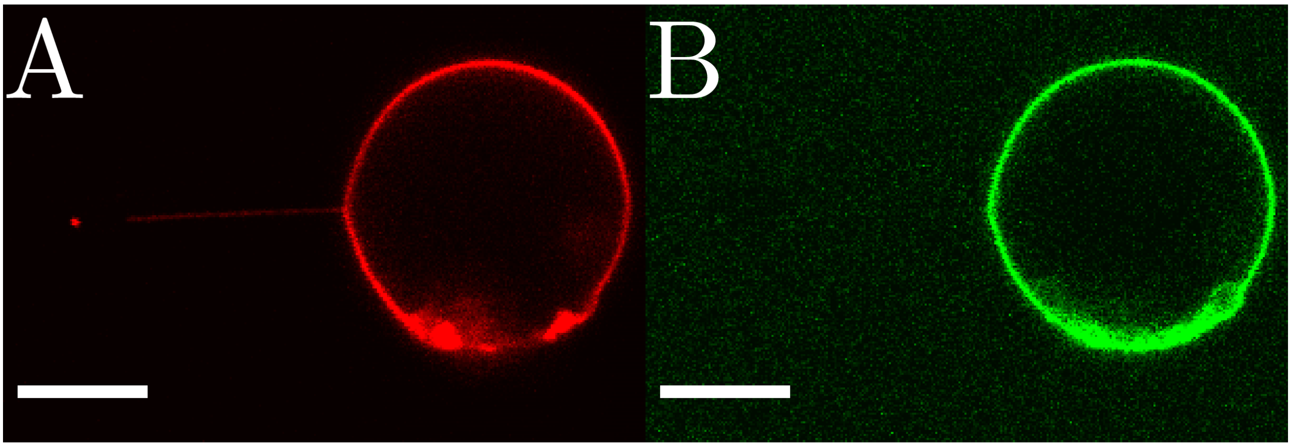


**Fig. S3.** Recombinant ANXA4 response to positive curvature. By adding recombinant proteins to the solution containing GPMVs labeled with vybrant DiD, we confirmed that an observation of binding of ANXA4 to positive curvature was not possible

| Laserpower | Slope [Intensity/concentration] |
|---|---|
| 20 | 25.63 |
| 30 | 51.10 |
| 40 | 77.59 |
| 50 | 100.99 |

**Supplementary Table 1.** Slopes from the plots in Supplementary Figure 1. The laser power was adjusted using acousto-optical tunable filter (AOTF). The calibration curves allow determination of the concentration of ANXA4-sfGFP based on the laser power used.
